# Reading of ingroup politicians’ smiles triggers smiling in the corner of one’s eyes

**DOI:** 10.1101/2023.08.11.553059

**Authors:** Edita Fino, Michela Menegatti, Alessio Avenanti, Monica Rubini

## Abstract

Capturing political support from spontaneous smile reactions detected in others’ faces can be used to gauge electorate preference. But will a smile elicited in the corner of one’s eye while reading of a favored politician smiling indicate positive disposition and political support for target candidates? From an embodied simulation perspective, we tested whether reading of an ingroup or outgroup politician smiling would trigger morphologically different smiles in faces of readers. In a reading task in the laboratory, participants were presented with subject-verb phrases describing left and right-wing politicians smiling or frowning while their facial muscular reactions were measured via electromyography (EMG) recording from the zygomaticus major (ZM, lip puller muscle), orbicularis oculi (OO, eye corner muscle) and the corrugator supercili (CS, wrinkler of the eyebrows). We expected and found that participants responded with a smile detected at the lip puller (ZM) and eye corner (OO) facial muscles when exposed to portrayals of smiling politicians of same political orientation, and reported more positive emotions towards these latter. When reading about outgroup politicians smiling, there was a weaker activation of the lip corner (ZM) muscle and no activation of the eye corner (OO) muscle, while emotions reported towards outgroup politicians were significantly more negative. Also, a more enhanced frown response (CS) was found for ingroup compared to outgroup politicians’ frown expressions. Present findings suggest that a politician’s smile may go a long way to influence electorates through both non-verbal and verbal pathways. They add another layer to our understanding of how language and social information shape embodied effects in a highly nuanced manner.

## 1. Introduction

In the political arena, people strive to understand how to capture political support (Menegatti & Rubini, 2013). At the nonverbal level, spontaneous facial expressions such as a fleeting smile detected in an unaware observer’s face, can be used to index affective reactions to political candidates and to reliably gauge electorate preference (McHugo & Lanzetta, 1991). An indication of positive affect in a smile can be inferred by the contraction of both the lip corner muscle (i.e., the zygomaticus major, ZM), which pulls the lips up into the familiar curve we most commonly identify with smiling, and the eye corner area (i.e., the orbicularis oculi, OO), also known as the Duchenne marker (Duchenne de Boulogne, 1876/1990; Ekman, Davidson, & Friesen, 1990.

Relevant to the present study is the evidence that unintended smile reactions are similarly triggered when seeing other people smiling, hearing them laughing, or reading about them smiling in the written media (Dimberg, 1990; Gallese & Lakoff, 2005; Magnée, Stekelenburg, Kemner & de Gelder, 2007; Niedenthal et al., 2009; Shaham, Mortillaro & Aviezer, 2020). This body of research indicates that as we perceive another person’s smile, we partially simulate or reproduce the smile in our sensorimotor system, in line with sensorimotor simulation accounts of facial expressions (Wood, Rychlowska, Korb & Niedenthal, 2016). Albeit automatic, facial effects triggered by perception of others’ expressions are modulated by social factors such as the similarity between observer and target, positive attitudes, social context and group membership (Hess, 2020; Hess & Fischer, 2014; Likowski, Mühlberger, Seibt, Pauli, & Weyers, 2008). Along these lines, the present study investigates a relatively neglected trigger of smiles, namely that of linguistic portrayals of politicians smiling. We employed facial electromyography (EMG) to detect smile responses during exposure to verbal depictions of political leaders smiling and frowning and examined whether different smiling patterns would emerge in faces of voters for ingroup but not outgroup politicians’ smiles reflecting positive affect and likely political support.

### 1.1 Smiles in the political context

Smiles are one of the main channels through which political leaders tend to induce a sense of happiness, confidence, and reassurance in others to the aim of increasing positive ratings, obtaining consensus and enlarging electoral support (Horiuchi, Komatsu & Nakaya, 2012; Masch, Gasner & Rosar, 2021). However, far from being simple readouts of positive affect, smiles have been shown to signal a wide range of socially relevant information and to regulate social interactions in a highly nuanced fashion (Hess, 2020; Niedenthal et al., 2010). Indeed, the political context offers one of the most ecologically valid settings for the smiles’ nuanced meanings to emerge (Stewart & Dowe, 2013). For instance, beyond genuine enjoyment, smiles are often used to signal absence of threat and a willingness to cooperate (i.e., affiliative smiles), as they may also be employed to express feelings of superiority and dominance towards a fierce opponent (i.e., dominance smiles) indicating in this case more negative rather than positive affect (Martin et al., 2018; Niedenthal et al., 2010; Rychlowska et al., 2017; Stewart, Bucy, & Mehu, 2015).

Smiles convey complex social information through the combined activation of a variety of facial muscles (Ekman & Rosenberg, 1997). For instance, when the contraction of the ZM which is commonly identified with a smile, is also associated with the contraction of the OO (the eye corner muscle responsible for creating small wrinkles around the eyes), then the smile is commonly perceived to express positive affect (Duchenne de Boulogne, 1876/1990; Ekman, Davidson, & Friesen, 1990). Hence, if a smile can be reach a person’s eyes, it is considered as displaying genuine enjoyment and as being more authentic (Ekman et al., 1990; Niedenthal et al., 2010; Sheldon et al., 2021; Stewart et al., 2010) compared to smiles activating the lip corner muscle only (ZM), which may often reflect social expression of politeness and may at times be also associated with negative affect (Ekman, 2009). This is especially relevant if smiles are displayed spontaneously, that is outside one’s conscious control, as related research suggests that when produced deliberately, Duchenne smiles might also be displayed in absence of positive feelings, (see Krumhuber & Kappas, 2022). However, spontaneous or not, it is generally agreed that Duchenne smiles lead to more favorable interpersonal perceptions. Indeed, people producing Duchenne smiles are rated more positively (for a meta-analysis see Gunnery & Ruben, 2016;) and they are facially responded to more than when producing non-Duchenne smiles (Hess & Bourgeois, 2010; Sims et al., 2012; Slessor et al., 2014).

### 1.3 Sensorimotor simulations accounts of facial effects

Research shows that smiles may trigger smiles not just through exposure to visual and auditory cues (Magnée et al., Gelder, 2006; Shaham et al., 2019), but also when one reads about someone smiling through a written text (Fino, Menegatti, Avenanti & Rubini, 2016; 2019; Foroni & Semin, 2009; Gallese & Lakoff, 2005; van Berkum, Struiksma1 & ‘t Hart, 2020). This is taken as further evidence that such facial effects reflect more complex processes than simply matched-motor, imitative responses in the strict sense (Hess & Fischer, 2013). Proponents of sensorimotor simulation accounts hold that as we perceive another person’s smile, we partially simulate or reproduce the smile in our sensorimotor system, which implicates the reactivation of related concepts, feelings and autonomic and behavioral changes (see Wood et al., 2016 for a review). This process enhances one’s capacity to respond emphatically (Blair, 2005) and recognize others’ emotions based on nuanced meanings of their facial expressions (Maringer, Krumhuber, Fischer, & Niedenthal, 2011; Niedenthal et al., 2010; Korb et al., 2014; Oberman, Winkielman, & Ramachandran, 2007; Rychlowska et al., 2014). Such claims are supported by evidence showing that when people are blocked in their ability to simulate smiles with their facial muscles (Borgomaneri et al., 2020; Foroni & Semin, 2009; Oberman et al., 2007), or when their face representation in the sensorimotor cortex is disrupted by brain stimulation (Paracampo, Pirruccio, Costa, Borgomaneri & Avenanti, 2018; Paracampo, Tidoni, Borgomaneri, De Pellegrino & Avenanti, 2017; Pitcher, Garrido, Walsh & Duchaine, 2008), their ability to correctly identify or judge emotional expressions of others is strongly impaired.

### 1.4 Social modulation of unintended facial effects

Although automatic, facial effects triggered by perception of others’ expressions are not impervious to social-contextual factors. Conceptual knowledge about the person doing the smiling and the social context in which the smile is displayed are important determinants of whether and how smiles will be corresponded, as they may affect observer’s expectations or inferences about the meaning expressed by smiles, reactivating related feelings and emotions accordingly (Calvo & Nummenmaa, 2016; Halberstadt et al.2009). Indeed, research has demonstrated rapid modulation by social information of such facial effects, suggesting a facilitation for people we favor and members of our own social group, and a reduction or at times suppression of the effect for people we dislike and outgroup members (for reviews, see Hess & Fischer, 2014; Seibt et al, 2015).

Indeed, while there is ample behavioral and neurophysiological evidence of stronger sensorimotor responding to ingroup than to outgroup members across a range of tasks, social groups and contexts (e.g., Avenanti, Surugu & Aglioti, 2010; Cazzato, Liuzza, Caprara, Macalusso & Alioti, 2015; Genschow & Schindler, 2016; Mondillon, Niedenthal & Droit-Volet, 2007; Rauchbauer, Majdandžić, Hummer, Windischberger, & Lamm, 2015; Riečanský, Kölble, Stieger, & Lamm, 2014), only a few studies have examined facial reactions to smiles in the political intergroup context. Early research employing electromyography to measure facial responses to politicians’ smiles (e.g., Bourgeois & Hess, 2008; McHugo & Lanzetta, 1991) has mainly relied on naturalistic stimuli such as short videoclips of politicians smiling. For instance, McHugo & Lanzetta (1991) presented participants with 30-second-long video excerpts of politicians smiling and found that the ZM response was more enhanced among supporters than opponents. These results were contrasted by a later adaptation of that study by Bourgeois and Hess (2008), who recorded facial activity at the ZM and OO while participants were exposed to 13s video sequences of smiling political leaders, finding that both ingroup and outgroup politicians’ smiles induced the same level of facial muscle activation.

A more recent approach (Fino, Menegatti, Avenanti & Rubini, 2019) focused on linguistic portrayals of left and right-wing politicians emotion expressions and recorded ZM and CS activity of left and right-wing voters. A more enhanced smile was evidenced amongst right-wing participants reaction in response to ingroup compared to outgroup politicians’ portrayals of smiles, whereas a similar facial response to ingroup and outgroup politicians’ smiles was evidenced among left-wing participants, who also showed with a higher ZM activity in response to outgroup politicians’ frowns, an indication of “*schadenfreude*” or pleasure derived by someone’s misfortune. This study was important in that it demonstrated that politicians’ smiles can induce smiles even when presented conceptually, through written media, extending previous research based on naturalistic stimuli. The use of linguistic material allows to control for the natural variability in the target politician’s smile type and intensity, which is inherent in research employing naturalistic stimuli (Bourgeois & Hess, 2008; McHugo & Lanzetta, 1991) and may independently affect observers’ behavior (Horiuchi et al., 2012). Notably, the favorable response detected in the smile reaction of right-wing participants to the left-wing political leader rising into power (Matteo Renzi) in that study, was a real-world demonstration that facial reactions to linguistic portrayals of political candidates can reliably predict electoral support, as they hinted at the massive support that Renzi would subsequently garner from both left-and-right wing electorates in the national elections following months later in Italy. However, these findings were limited by the over reliance on ZM activity as the only index of readers’ smiling reaction and such one-size-fits-all approach to smile reactions did not allow for disambiguation of more nuanced smiling patterns amongst supporters and opponents. For instance, it cannot be ruled out that beyond a similar ZM activation, left-wing participants may have been reacting with different kind of smiles altogether, reserving a more malicious one (i.e., dominant smile) for outgroup politicians as suggested by their *schadenfreude* response.

### 1.5 The present study

Current evidence about facial reactions to politicians’ smiles is mixed, with some studies reporting that smiles are enhanced for ingroup compared to outgroup leaders (McHugo et al., 1998), and other studies showing a similar response across political groups (Bourgeous & Hess, 2008; Fino et al., 2019 - left-wing participants). This begs the question of whether there may be more fine-grained differences underlying such effects. Prior work has employed different experimental procedures and was focused mainly on the ZM activation (except for Bourgeois & Hess, 2008). The question of whether the activation of the OO – a key facial muscle involved in smiles signaling positive affect (otherwise known as Duchenne smiles) – could further qualify facial responses to politicians’ smiles, depending on political affiliations, deserves further attention. To date, no study has examined whether the Duchenne marker, indicating positive affect in a smile, can be observed when participants are facially responding to linguistic representations of others’ smiling, as most research on Duchenne smiles has traditionally relied on visual stimuli. The present work addresses these gaps by investigating whether reading of ingroup or outgroup politicians’ portrayals of smiles induces qualitatively different smile reactions among readers of same and opposing political orientation, reflecting enjoyment for ingroup but not outgroup politician’s smile depictions.

Differently from experimental procedures employed in previous research on Duchenne smiles often relying in deliberately produced facial expressions (Kruhumber & Kappas, 2022), we measured unintended facial reactions to linguistically presented stimuli. Moreover, by indexing facial responses across different time windows in a 0 - 3s post-stimulus onset interval, this is also the first study to explore the temporal dynamic of facial reactions to verbal descriptions of politicians’ smiles. In a laboratory reading task, participants were exposed to verbal phrases describing ingroup and outgroup politicians smiling and frowning, while their spontaneous facial reactions were measured via EMG from the ZM, OO and the Corrugator supercili (CS, wrinkler of the eyebrows). Based on previous evidence showing mixed results in terms of ZM discriminating between ingroup and outgroup smiles, we expected that facial responses to phrases depicting smiling ingroup politicians would involve a significant activation in both the lip puller (ZM) and eye corner (OO) facial muscles, indicating positive affect, which would be further corroborated by higher levels of positive emotions reported for ingroup compared to outgroup politicians. In response to outgroup politicians’ smiles, we expected to find lesser or similar activation of the ZM and no activation of the OO, as well as higher levels of negative emotions reported towards these later. In addition, we expected a more enhanced frown response (CS) for ingroup compared to outgroup politicians’ frown expressions.

## 2. Materials and Methods

### 2.1 Participants

Thirty undergraduate students at the University of Bologna (27 females, mean age 22 years old) participated in the study for academic credit. Data were gathered from May to October 2013 at the University of Bologna. Participants were selected from a pool of 120 students filling out a political identification scale from 1 (left) to 7 (right). Those agreeing to participate in the present study reported being left-wing and scored below 3 on the political identification scale. All subjects had normal or corrected-to-normal vision, were right-handed and were naïve to the purpose of the experiment. The study was not preregistered, the protocol was approved by the Bioethics committee of the University of Bologna and was carried out in accordance with the ethical standards of the 2013 Declaration of Helsinki (World Medical Association, 2013). Prior to the start of the experiment participants read the ethical approval statement and provided written informed consent. Data from one subject were excluded from final analysis due to technical problems with electrode adherence.

### 2.2 Stimuli and Procedure

#### 2.2.1 Behavioral task and general procedure

Participants sat approximately 60 cm from a 19-inch computer monitor. They performed a judgement task requiring them to read a series of subject-verb phrases sequentially presented on the center of the monitor and rate them on a 5-point liking scale (1 = *I don’t like it at all*; 5 = *I like it very much*). Each sentence was preceded by a fixation cross lasting 1s, which was followed by the target phrase remaining on the screen for 3 s and followed by the liking scale that disappeared upon response (mean RT was ∼ 4 s). Then the screen remained blank for 1s in the inter-trial interval. Thus, the inter-stimulus interval was ∼10 s allowing sufficient time for facial muscles to relax after stimulus presentation.

The stimulus material consisted in 12 Italian verbs (6 positive and 6 negative emotion expressions) and 6 neutral fillers taken from Fino et al. (2019). Positive verbs included: *to smile* (sorridere), *to laugh* (ridere), *to enjoy* (gioire); Negative verbs included: *to frown* (corrucciare), *to grin* (aggrotare), *to get angry* (arrabbiarsi). Fillers included: *to work* (lavorare); to walk (camminare). All verbs were matched for word length and frequency (see masked). Verbs were presented in the present tense and were embedded into subject-verb sentences with each target verb being attributed to either a left-or right-wing politician (e.g., ‘*Bersani smiles*’, ‘*Alfano frowns’*). Based on a pilot study [masked], we selected two left-wing politicians (Bersani and Renzi) and two right-wing politicians (Alfano and Berlusconi) from the main political parties (Democratic party and People of Freedom, respectively) at the time of the study. For each participant three blocks of stimuli were presented with each block containing all the 12 emotion expression verbs and 6 neutral verbs attributed to 4 left-and right-wing politicians, resulting in 72 stimuli presentations per block, and a total of 216 trials altogether. Prior to the start of the experiment, 10 practice trials were administered for participants to familiarize themselves with the computer task. During the experiment, the verbal stimuli were presented in a random order with E-prime software (Psychology Software Tools, Pittsburgh, PA).

After the task, participants completed a questionnaire including manipulation checks and assessing their attitudes on a multiplicity of measures. They were first asked to indicate their political orientation on the same item used prior to the experiment and report on the valence of each stimulus sentence (see next paragraph). Then they indicated the political affiliation of each politician by choosing between two options (left or right) and rated how much each target politician represented the political wing they had indicated. In the end participants were asked about their ideas regarding the purpose of the experiment, they were debriefed and dismissed. None of the participants was aware of the study hypotheses and none suspected that facial muscular reactions were being measured.

#### 2.2.2 Apparatus and Data acquisition

Facial muscle activity was measured during stimulus presentation using miniature Ag/AgCl surface electrodes (4mm) attached over the left ZM, OO and CS muscles. To conceal the real purpose of the study participants were initially told they were participating in a study on language of politics that involved a reading task during which skin conductance levels would be recorded via sensors placed on the face. Site preparation and electrode placement were done following standard procedure guidelines (Fridlund & Cacioppo, 1986; Tassinary & Cacioppo, 2000). The skin was cleaned and prepared for electrode placement to reduce skin impedance to less than 10 kΩ. The raw EMG activity was measured with Biopack Systems MP36 data acquisition unit at a sampling rate of 1000 Hz using two bipolar channels and a gain of 1,000. The EMG signal was pass filtered with a 20-250 Hz passband and a 50 Hz notch filter and was full wave rectified offline.

### 2.3 Dependent variables

#### 2.3.1 Manipulation checks

To check participants’ political orientation, they were asked to report on the item: “Where would you locate yourself politically in a continuous scale from 1(*left*) to 7 (*right*)”. To check whether the target politicians were considered as being left- or right-wing, participants were asked how much they represented the right- and the left-wing respectively on a 7-point Likert scale from 1 *(not at all)* to 7 *(very much)*. To check whether positive and negative subject-verb sentences were evaluated as positive and negative respectively, participants were asked to assess the valence of each phrase on a 7-point Likert scale from 1 (*very positive*) to 7 (*very negative*).

#### 2.3.2 Facial muscle EMG activity

Smile reaction patterns were assessed through EMG recording at the ZM and OO muscle sites. Frown reactions were measured by EMG recording of the CS (brow wrinkler muscle) activity. The EMG signal in the time window of interest (1000ms pre-stimulus to 3000ms post-stimulus onset) was rectified and root mean square (RMS) transformed. On each trial, mean EMG amplitude in the post-stimulus period was baseline-corrected using mean EMG amplitude during the pre-stimulus period. Phasic facial EMG responses (in microvolts, mV) were scored across bins of 100ms and averaged over intervals of 500ms during the 3s of stimulus presentation. The EMG signal was visually inspected offline and screened for electrical noise and artifacts caused by muscle movements and blinks, resulting in approximately 5% of 100ms intervals of EMG data and trials being excluded from the analysis. Less than 2% of trials were excluded from the analyses using the standard deviation (SD) method (Wilcox, 1992) with the criterion value standing at 3 SDs per muscle. Before statistical analysis, EMG data on each facial muscle were averaged over 18 trials with the same emotional expression for left-and right-wing politicians.

#### 2.3.3 Evaluative rating

To test whether politicians’ facial expressions described in the subject-verb sentences elicited evaluative judgments in the reader, we asked participants to rate each sentence on a liking scale. Liking ratings of subject-verb sentences were provided after stimulus presentation by right-hand clicking on a 5-point Likert scale from 1 (*I don’t like it at all*) to 5 (*I like it very much*).

#### 2.3.4 Emotions towards target politicians

Participants’ emotional attitudes towards target politicians were assessed through a PANAS scale consisting of 3 positive (joy; enthusiasm; excitement) and 3 negative (anger; sadness; disappointment) emotions. The respondents indicated the degree to which thinking about the target politician elicited the given emotion on a 7-point Likert scale from 1 (*not at all*) to 7 (*very much*).

#### 2.3.5 Voting intention towards target politicians

To assess participants’ voting intentions we asked them to report on the likelihood of their voting for the target politicians in the coming elections on a 7-point Likert scale from 1 (*not at all likely*) to 7 (*very likely*).

## 3. Results

Electrophysiological and behavioral data were inspected for normality and then analyzed using a series of repeated measure analysis of variance (ANOVA) and correlational analysis, as detailed below. Post-hoc analysis was conducted using Bonferroni correction for multiple comparisons. Partial eta squared (η_p_2) was computed as a measure of effect size for significant main effects and interactions. By convention, η_p_2 effect sizes of ∼ .01, ∼.06, and ∼.14 are considered small, medium and large, respectively (Cohen J, 1992).

### 3.1 Manipulation check

All participants reported being left-wing (*M*=1.96, *SD*=0.56). Alfano (*M*=5.24; *SD*=1.02) and Berlusconi (*M*=5.65; *SD*=0.93), were considered as representing the right-wing, whereas Bersani (M=4.92; SD=1.16) and Renzi (M=4.80; SD=1.20) were considered as representative of the left-wing, respectively. Repeated measures ANOVA showed that positive emotion verbs were considered as more positive (M=5.68; SD=0.59) than negative emotion verbs (M=2.52; SD=0.56), (*F*_1,28_=334.901, *p*<.0001, η_p_^2^=.923).

### 3.2 Facial muscle activation pattern in response to ingroup and outgroup politicians’ expressions

EMG data were analyzed using a 2 (Political orientation: left-wing/ingroup, right-wing/outgroup) × 2 (Facial expression: smile, frown) × 6 (Time: 1-6 bins of 500ms) repeated measures ANOVA for the ZM, OO and CS muscles separately.

#### 3.2.1 Zygomaticus Major (ZM)

The Political orientation × Facial expression × Time ANOVA on ZM data showed a main effect of Political orientation (*F*_1,28_=5.877, *p*=.022, η_p_^2^=.173). Facial muscle activation was higher for ingroup (*M*=.013, *SE*=.02), compared to outgroup politician’s expressions (*M*=-.046, *SE*=.03). The two-way Political orientation × Facial expression interaction was significant (*F*_1,28_=11.712, *p*=.002, η_p_^2^=.295), which was further qualified by the significant Political orientation × Facial expression × Time interaction (*F*_5,140_=7.501, *p*=.002, η_p_^2^=.211). Figure 1 shows a consistent and slowly increasing ZM activation when participants read about smile expressions of ingroup politicians, and opposite trends when reading about ingroup frown expressions or when reading about outgroup smiles, with modulation peaking in the last time bins of the explored time window (2000-3000ms). Pairwise comparisons (based on Bonferroni tests) revealed that ZM activation was significantly higher when participants read smile expressions attributed to ingroup compared to outgroup politicians across all time bins (0 – 3000ms; all *p*s<.011; Fig.1). Moreover, while reading frown expressions, there was a suppression of ZM activity for frowns attributed to ingroup compared to outgroup politicians (Fig.1), with significant differences emerging only at bin 1 (p=.033), and bin 6 (p=.035). Additionally, when participants were presented with smile and frown expressions of ingroup politicians their ZM activation was significantly higher for the former compared to the latter across all time bins (all *p*s<.002). Instead, there was no consistent difference in the ZM activation for smile and frown expressions of outgroup politicians (all *p*s>.08).

**Figure 1.**
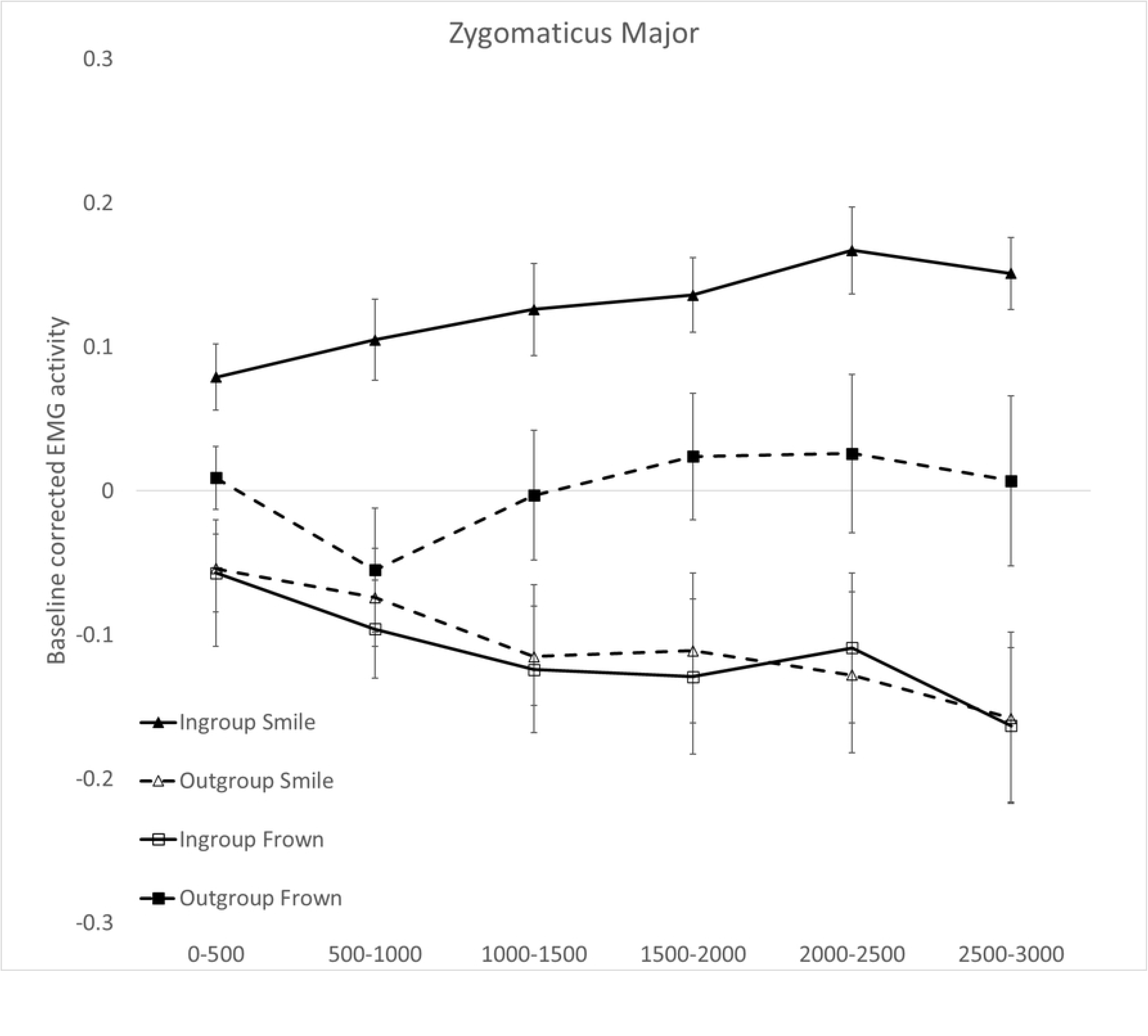
Zygomaticus Major muscle activity for left-wing versus right-wing politicians’ verbal expressions of smiles (panel A) and frowns (panel B). Responses are broken down in six 500ms-time windows of stimulus exposure (0 to 500ms; 500 to 1,000ms; 1,000 to 1,500ms; 1,500 to 2,000ms; 2,000 to 2,500ms; 2,5000 to 3,000ms). Error bars indicate standard errors of the mean.

#### 3.2.2 Orbicularis Oculi (OO)

The Political orientation × Facial expression × Time ANOVA on OO data showed a main effect of Political orientation (*F*_1,28_=15.228, *p*=.001, η_p_^2^=.352) with higher EMG activation for ingroup (*M*=.110, *SE*=.02) compared to outgroup politician’s expressions (*M*=-.034, *SE*=.04); a main effect of Facial expression (*F*_1,28_=10.803, *p*=.003, η_p_^2^=.278) with higher EMG for positive (*M*=.107, *SE*=.02) compared to negative (*M*=-.031, *SE*=.04) facial expressions; and a main effect of Time (*F*_5,140_=17.205, *p*<.0001, η_p_^2^=.381), with higher activations in bin 1 (0-500ms, M=0.111mV, SE=0.03), bin 2 (500–1000ms, M=0.158mV, SE = 0.04) and bin 3 (1000-1500ms, M= 0.100mV, SE = 0.03) compared to bin 4 (2000ms, M= -0.004mV, SE = 0.03), bin 5 (2500 ms, M= -0.047mV, SE = 0.03) and bin 6 (3000ms, M= -0.095mV, SE = 0.04; all *p*s < 0.0001).

The Political orientation × Facial expression interaction was also significant (*F*_1,28_=46.925, *p*<.0001, η_p_^2^=.635) and it was further qualified by the significant Political orientation × Facial expression × Time interaction (*F*_5,140_=6.577, *p*<.0001, η_p_^2^=.196). Figure 2 shows a strong OO activation when reading about smile expressions attributed to ingroup politicians with an early peak around the second time bin (500-1000ms), and a weaker comparably early OO activation when reading about outgroup frown expressions. Pairwise comparisons (based on Bonferroni tests) revealed that participants reacted with a higher OO activation when reading of smile expressions attributed to ingroup compared to outgroup politicians, across all time bins (0–3000ms upon stimulus onset; all *p*s<.0001; Fig.2). When participants were presented with frowns, OO activation was significantly suppressed for ingroup compared to outgroup politicians’ expressions, across time bins (all *p*s<.010; Fig.2). In addition, smiles compared to frown expressions of ingroup politicians were associated with greater OO muscle activation in all time bins (all *p*s<.0001), whereas the inverse pattern was observed for expressions of outgroup politicians: smiles induced significantly lower activation of the OO muscle compared to frowns, across all time bins (all *p*s<.050).

**Figure 2.**
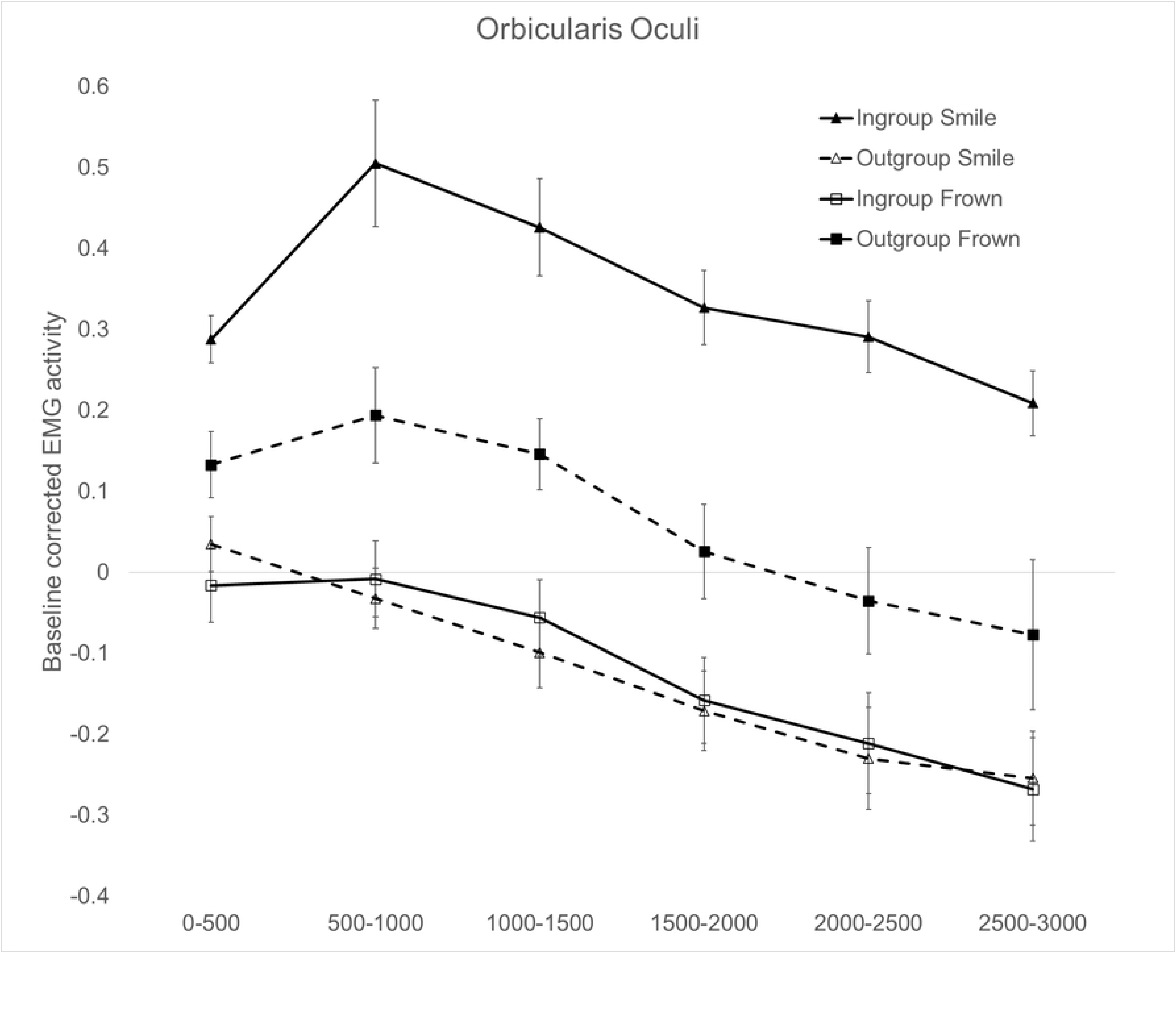
Orbicularis oculi muscle activity for left-wing versus right-wing politicians’ verbal expressions of smiles (panel A) and frowns (panel B). Responses are broken down in six 500ms-time windows of stimulus exposure (0 to 500ms; 500 to 1,000ms; 1,000 to 1,500ms; 1,500 to 2,000ms; 2,000 to 2,500ms; 2,5000 to 3,000ms). Error bars indicate standard errors of the mean.

#### 3.2.3 Corrugator Supercili (CS)

The Political orientation × Facial expression × Time ANOVA on EMG data recorded on the corrugator supercilii showed a main effect of Political orientation (*F*_1,28_=30.159, *p*<.0001, η_p_^2^=.519), with higher CS activity for ingroup (*M*=.137, *SE*=.03) compared to outgroup politician’s expressions (*M*=-.014, *SE*=.02); a main effect of Facial expression (*F*_1,28_=13.829, *p*=.001, η_p_^2^=.331), with higher activation for negative (*M*=.135, *SE*=0.03) compared to positive (*M*=-.012, *SE*=0.03) facial expressions; and a main effect of Time (*F*_5,140_=13.864, *p*=.019, η_p_^2^=.121), with higher CS activation during bin 1 (0-500ms, M=.105, SE=.02) and bin 2 (500–1000ms, M=.117, SE=.03) compared to bin 3-6 (1000-3000ms, M=.063-0.14, SE=.03-.04; all *p*s<.030).

The Political orientation × Facial expression interaction (*F*_1,28_=41.138, *p*<.0001, η_p_^2^=.595) was also significant which was further qualified by the significant Political orientation × Facial expression × Time interaction (*F*_5,140_=10.937, *p*<.0001, η_p_^2^=.281). Figure 3 shows strong sustained CS activity when reading frown expressions of ingroup politicians with activation peaks at bin 5 (2000-2500ms) and a weaker late deactivation when reading about smile expressions of ingroup politicians. Pairwise comparisons (based on Bonferroni tests) revealed that the activation of the CS was higher when participants read about frown expressions of ingroup compared to outgroup politicians across all time bins (all *p*s<.0001; Fig.3). In terms of smile expressions, a greater suppression of the CS activation was evidenced for expressions of ingroup compared to outgroup politicians (Fig.3), with significant differences emerging at bin 1 (p=.027), bin 5 (p=.023) and bin 6 (p=.027). When participants were presented with expressions of ingroup expressions their CS response was higher for frowns compared to smiles in all time bins (all *p*s<.0001). Instead, the reverse pattern was observed when they were reading expressions of outgroup politicians: CS activation was lower for frown compared to smile expressions, with significant differences emerging at bin 1 (0-500ms, p=.030) and bin 3 (1000-1500ms, p=.006).

**Figure 3.**
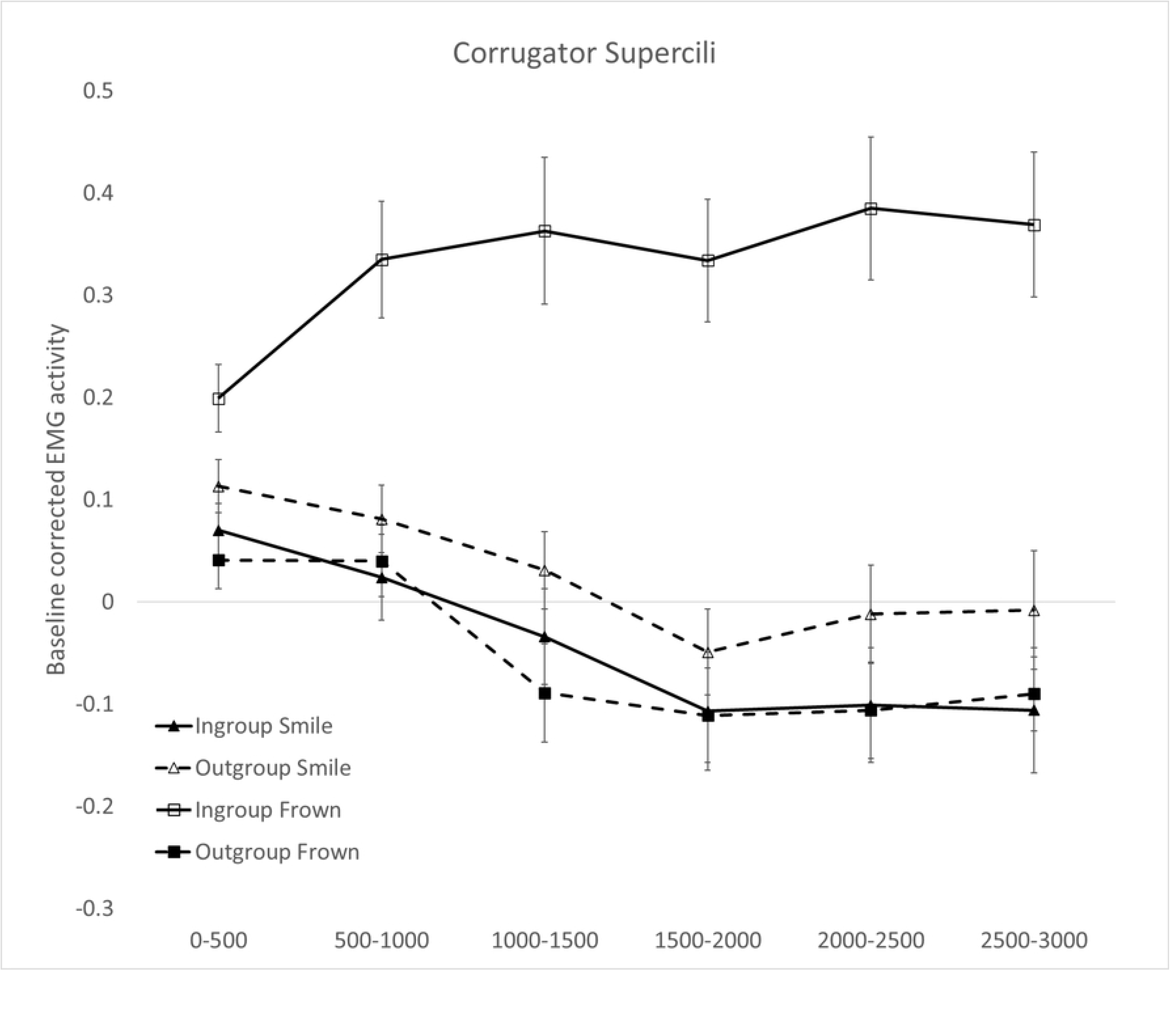
Corrugator Supercilii muscle activity for left-wing versus right-wing politicians’ verbal expressions of frowns (panel A) and smiles (panel B). Responses are broken down in six 500ms-time windows of stimulus exposure (0 to 500ms; 500 to 1,000ms; 1,000 to 1,500ms; 1,500 to 2,000ms; 2,000 to 2,500ms; 2,5000 to 3,000ms). Error bars indicate standard errors of the mean.

### 3.3 Evaluative ratings

To test whether participants liked to a different extent the facial expressions displayed by politicians of their own compared to the opposite political orientation, self-report data were analyzed in a 2 (Facial expression: smile, frown) × 4 (Politicians: Alfano, Berlusconi, Bersani, Renzi) repeated measures ANOVA. Results revealed the main effects of Politician, (*F*_3,84_=34.455, *p*<.0001, η_p_^2^=.552) indicating higher liking rates for ingroup compared to outgroup politicians (see Table 1). This was further qualified by the significant interaction between Politician and Facial expression (*F*_3,84_=33.283, p<.0001, η_p_^2^=.543). Pairwise comparisons (based on Bonferroni tests) showed that participants reported significantly higher liking rates for the smiling of ingroup compared to outgroup politicians Berlusconi (all *p*s<.001). Instead, similar rates of liking were reported for frowning of ingroup compared to outgroup politicians Berlusconi (all *p*s>.480).

**Table 1.** Mean (± SD) scores on liking of politician verbal expressions reported by participants for each target political leader.

| Liking of politician verbal expressions |  |  |  |  |
| --- | --- | --- | --- | --- |
|  | Target Politicians | All emotion expressions | Positive emotion expressions | Negative emotion expressions |
| Left-Wing (ingroup) | Bersani | 3.02 ± 0.08 | 3.46 ± 0.71 | 2.57 ± 0.45 |
| | Renzi | 2.78 ± 0.07 | 3.37 ± 0.71 | 2.53 ± 0.36 |
| Right-Wing (outgroup) | Alfano | 2.26 ± 0.10 | 1.95 ± 0.67 | 2.61 ± 0.72 |
| | Berlusconi | 2.17 ± 0.11 | 1.73 ± 0.88 | 2.65 ± 0.86 |

### 3.4 Emotions toward target politicians

Affective responses were analyzed into a 2 (Emotions: positive, negative) × 4 (Politician: Alfano, Berlusconi, Bersani, Renzi) repeated measures ANOVA. The analysis showed a main effect of Emotion (*F*_1,19_=28.045, *p*<.0001, η_p_^2^=.596) showing higher scores for negative (*M*=3.64, *SE*=.14) relative to positive emotions (*M*=2.69, *SE*=.11); a main effect of Politician, (*F*(3, 57) = 9.864, *p* < .0001, η_p_^2^ = .342) showing different scores across target politicians; and a significant interaction between Emotion and Politician (*F*_3,57_=102.085, *p*<.0001, η_p_^2^=.843; Fig. 4). Pairwise comparisons (based on Bonferroni tests) showed that participants reported more positive emotions towards ingroup politicians Bersani (*M*=4.16, *SD*=1.03) and Renzi (*M*=4.14, *SD*=1.55) compared to outgroup politicians Berlusconi (*M*=1.19, *SD*=0.40; all *p*s<.001) and Alfano (*M*=1.27, *SD*=0.38; all *p*s<.001). Additionally, participants reported more negative emotions towards outgroup politicians Berlusconi (*M*=5.95, *SD*=1.03) and Alfano (*M*=4.21, *SD*=0.96) compared to ingroup politicians Bersani (*M*=2.35, *SD*=0.97; all *p*s<.001) and Renzi (*M*=2.05; *SD*=.99; all *p*s<.001).

**Figure 4.**
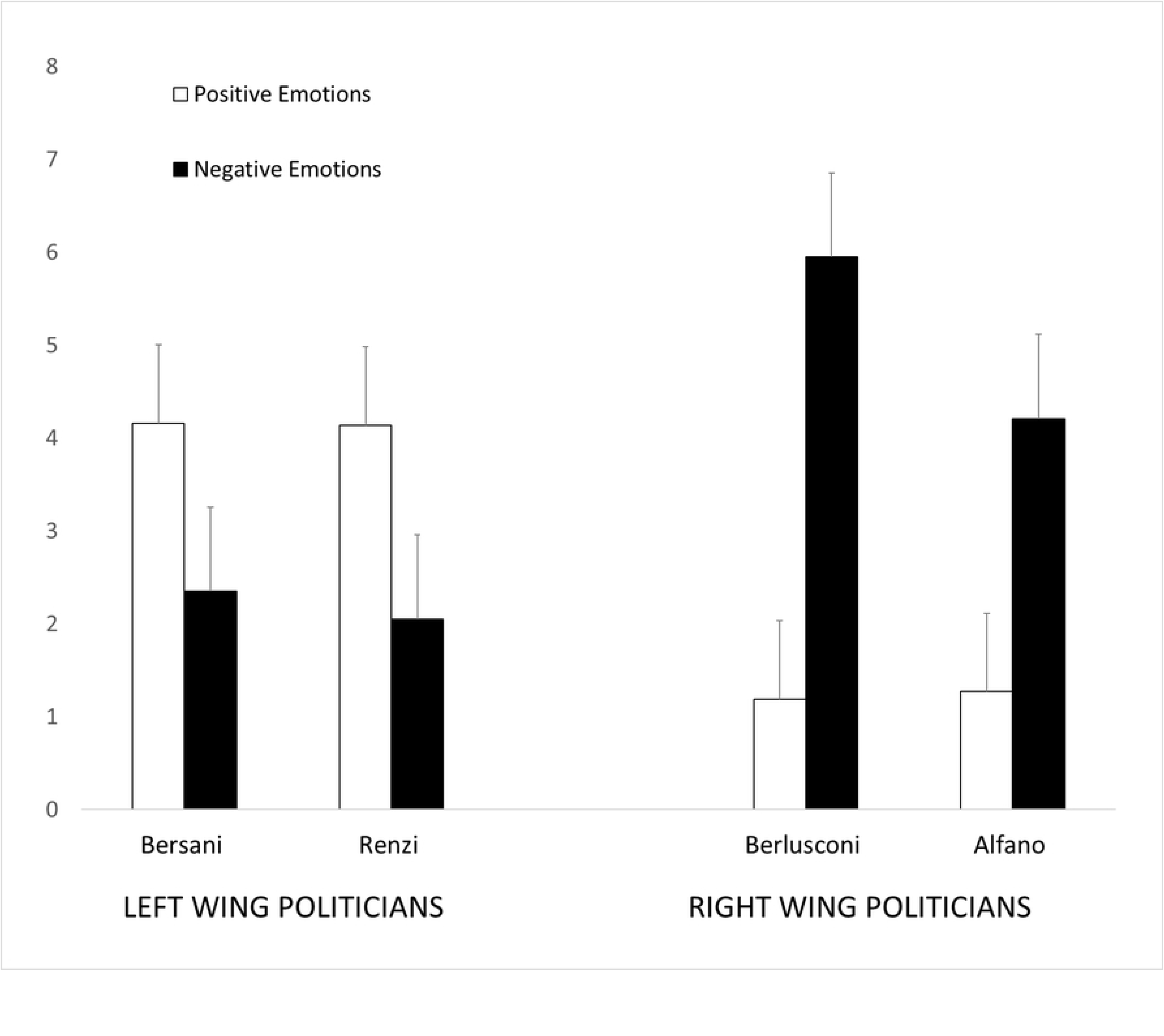
Positive and negative emotions reported for ingroup (left-wing) and outgroup (right-wing) target politicians. Error bars indicate standard errors of the mean. * *p <* .05. ** *p <* .01. *** *p <* .001.

### 3.5 Voting Intention towards target politicians

The ANOVA performed on voting intentions ratings showed that participants would most likely vote for ingroup politicians Bersani (M=5.10, SE=.34) and Renzi (M=4.63, SE=.30), compared to outgroup politicians Alfano (M=1.05, SE=.05) and Berlusconi (M=1.15, SE=.07) in the upcoming elections (*F*_3,84_=91.960, *p*<.001, η_p_^2^=.767).

### 3.6 Relationship between facial responses, evaluative judgements, emotions and voting intentions towards political leaders

Finally, we examined correlations between facial EMG responses to ingroup and outgroup facial expressions, evaluative judgments, reported emotions and voting intentions towards ingroup and outgroup politicians. We did not find significant correlations between EMG and behavioral data.

## 4. Discussion

Politicians may strategically use smiles to influence their voters’ behavior (Horiuchi et al., 2012; Masch et al., 2021; Sullivan & Masters, 1988), but will reading about a politicians’ smile spontaneously trigger nuanced smile reactions, which may differ depending on whether readers share the same or opposing political orientation? In the present study we examined facial EMG activity to test whether reading about ingroup or outgroup politicians’ smiling would elicit morphologically different smiles in the faces of readers. As expected, we found that participants facially responded with a smile pattern involving the activation of both lip puller (ZM) and eye corner (OO) facial muscles to phrases describing ingroup but not outgroup politicians smiling. By going beyond a one-size fits-all approach to detecting smiles, and by examining the temporal dynamic of facial muscle activation we were able to capture a more nuanced smile reaction in our participants revealing fine-grained differences in early responses to ingroup compared to outgroup politicians’ smiling portrayals. This is the first study to show that a smile pattern which is underpinned by the activation of both lip puller (ZM) and eye corner (OO) facial muscles, can be observed not just when one sees a loved political leader smiling, but also when reading about their smiling expressions in written media. Interestingly while in all muscles we were able to detect emotionally congruent reactions to ingroup politicians’ expressions (i.e., ZM and OO activation for ingroup smiles and CS activation for ingroup frowns), the OO muscle showed the earliest peak of activation and appeared particularly sensitive to the ingroup vs. outgroup manipulation as compared to the other muscles.

These findings importantly extend previous research based on exposure to visual (Bourgeois & Hess, 2008; McHugo & Lanzetta, 1991) or auditory cues (Magnée et al., 2006; Shaham et al., 2019) as they are the first to suggest that, with perceptual features of target’s smile stringently controlled through the use of linguistic material, mere conceptual knowledge about the target is able to modulate the readers’ facial response in a highly nuanced fashion, so that a smile is differently performed through the recruitment of one’s facial muscles when reading about smiles of ingroup vs. outgroup politicians (Niedenthal et al., 2010). Results extend previous findings of Fino et al., (2019) by showing that there can be fine-grain distinctions in the readers’ smile response, when the eye corner (OO) muscle activation is concurrently tracked with that of the lip corner muscle (ZM). Not all smiles are equal they say, and when our participants were reading of ingroup politicians smiling, a spontaneous contraction could be captured at the level of the eye corner muscle (OO) in concurrence with that of the lip puller muscle (ZM), suggesting a Duchenne (aka positive affect) smile reaction for ingroup but not outgroup politicians’ smiles (Ekman et al., 1990; Martin et al., 2017). This was confirmed by the reported emotions towards target politicians which were significantly more positive for ingroup and more negative for outgroup political leaders, in line with our predictions.

By highlighting the importance of the eyes in detecting highly nuanced smile reactions (Niedenthal et al., 2010), our findings further extend previous research focusing on the Italian political context showing that a politician’s eyes are highly salient and grab the attention of voters automatically. In their study, Liuzza, and colleagues (2011) examined how the eye gaze of a prominent right-wing Italian politician (Berlusconi) had a strong catching power over the attention of potential voters, potentiating gaze following behavior of supporters and inhibiting that of opponents. By using a different experimental paradigm, in the present study we measured reflexive smile patterns emerging very rapidly in the eye corner area of supporters, demonstrating a highly differentiated smile response to ingroup but not outgroup smile displays. What is remarkable is that such fine-grained differences in supporters’ smile reactions were observed as they facially responded to language portraying politicians smiling. The pivotal role of language as a means of exerting influence for the achievement of individual and social and political goals has been widely demonstrated (e.g., Menegatti & Rubini, 2013; Rubini, Menegatti & Moscatelli, 2015). Our findings go a step further into deepening our understanding of how language can sensibly shape one’s unintended behavior through seamlessly nuanced embodied effects.

The fact that the facial reaction pattern underpinned by the activation of both ZM and OO muscles - which reflects the signature of Duchenne smiles – was evidenced only for linguistic portrayals of ingroup but not outgroup politicians’ smiles suggests that expressions of ingroup and outgroup leaders are differently processed. This is consistent with research on social modulation of facial effects demonstrating that emotionally congruent facial reactions are tendentially more enhanced for ingroup members, people we like and those towards whom we have affiliative intentions (Hess & Blairy, 2001; Likowski et al., 2008; Steel et al., 2010). Our findings also converge with evidence showing that Duchenne smiles are associated with greater liking and perceived psychological proximity (Bogodistov & Dost, 2017), which is expectedly higher towards ingroup compared to outgroup members. In line with established research lines on intergroup perception (Cikara & Van Bavel, 2014; Han, 2018; Hewstone, Rubin & Willis, 2002), people show positive bias towards ingroup members as compared to outgroup members; therefore, they tend to judge an ingroup smile as more genuine and authentic or as signifying positive feelings or intentions. Conversely, in the case of outgroup smiles, it is more probable that the perceivers believe that the smile masks a feeling of superiority, a desire to manipulate or lie, or simply negative affect, which are all the more expected in the political context (Stewart, Senior & Bucy, 2012). Hence, in absence of more contextual information, it may be that while reading about ingroup and outgroup politicians’ smiles participants may have made inferences about smiles meaning and/or genuineness, which may have been reflected in their different smile response. This interpretation is consistent with the pattern of reported liking rates by our participants which were significantly higher for ingroup and lower for outgroup politicians’ smiles. As suggested by previous research, smiles that are perceived as more authentic and truer are liked more and elicit more positive emotions compared to smiles that are perceived as fake or as expressing superiority, which are associated with more negative emotions (Ekman, 2009; Ekman et al., 1990; Niedenthal et al., 2010; Stewart et al., 2010). Although it has been noted that Duchenne smiles can also be displayed deliberately and in absence of positive affect (Kruhumber & Kappas, 2022), the fact that we measured spontaneous facial reactions while participants were unaware that their facial muscle activity was being recorded rules out the possibility that they may have intentionally smiled with their eyes.

The present research goes beyond the one-size-fits-all approach to smile responses which assumes ZM activation to be the main and only index of smile reactions. Indeed, we found that not only the ZM but also the OO muscle appeared consistently engaged in the task and sensitive to our language manipulations, and showed the highest and earliest peaks of EMG activations relative to the other muscles. By the combined measurement of more facial muscle activity parameters, based on the evidence on smile variability (Martin et al., 2017; Niedenthal et al., 2010), and the analysis of the temporal dynamic of facial muscle activation we were able to reach a more in-depth analysis of differences in facial reactions, which may have been previously masked by measurement approaches employed in other studies. The fact that we found significant ZM and OO activation (reflecting the Duchenne marker) in response to ingroup but not outgroup smiles, opens the way for a different interpretation of what has up to now been considered an intriguing empirical evidence in this area, namely that smiles are sometimes indiscriminately corresponded across social groups and contexts (Hess, 2021). While smiles are often used to smooth social interactions (i.e., affiliative smiles), previous research may have neglected more subtle differences in smile reactions. In relation to politicians’ smiles, the only previous study to concurrently assess responses at the ZM and OO muscles found no evidence for increased activity in OO for ingroup relative to outgroup politicians (Bourgeois & Hess, 2008). However, it should be noted that this study employed ecological dynamic stimuli and assessed facial reactions over an extended 13-s time frame. The question of whether stimuli related features or timeframe exposure may have influenced results cannot be completely ruled out as contractions of OO might be sensitive to the time of post-stimulus onset with activity peaking in earlier windows and diminishing with time as showed by previous research (Slessor et al., 2014). Hence, the importance of a temporal dynamics analysis which may reveal more subtle differences in facial muscle activation as it enfolds through time.

However, our research is not exempt from limitations. For one thing, ours was a single study involving a sample of left-wing participants. More studies are required to further substantiate our results with right-wing participants as well, ruling out potential confounding factors related to political ideology that may have influenced our results. Furthermore, our sample was predominantly female with target politicians examined being all male, hence we were unable to examine gender related effects. Recent studies have shown that female politicians tendentially display more intense and affiliative type of smiles compared to their male counterparts and are judged more negatively if they don’t (Senior et al., 2019). More studies are needed to examine the modulation effect of gender in whether and how politicians’ smiles are perceived and facially responded to. Additionally, we did not provide information about the interaction context in which politicians’ smiles were displayed, whether they were in response to a praising comment (i.e., enjoyment smile) or an angry opponent’s remark (i.e., dominant smile). Future studies should do so and consider a more stringent manipulation of the meaning of the smiles presented through verbal material. For instance, presenting phrases such as “Politician smiles joyfully” vs. “Politician smiles politely” or with superiority, may be a useful approach to examine fine grain differences in supporters’ and opponents’ smile reaction patterns.

Despite these limitations our findings are the first to highlight how language and social information can shape embodied effects in a highly differentiated manner, which can be of relevance to the field of social and political communication. They also offer some evidence suggesting that existing differences in empirical findings on facial reactions to smiles in the intergroup context might be reconciled by going beyond one-size-fits-all approaches that rely on assessment of ZM activity only. This has implications for future research focusing on disambiguating the facial response to smiles, whether presented conceptually or not, in different social and emotional contexts. Integrating more facial markers involved in different smile expressions would also allow to capture more complex and nuanced smile patterns. Moreover, using linguistic portrayals of others’ smiling offers a useful approach in terms of more stringently testing social interaction and contextual factors. Also combining additional neuro/psycho-physiological indicators of emotional states while facially responding to different types of smiles, as suggested by recent research (Martin et al., 2018; Rychlovska et al., 2017), may provide a more comprehensive picture of the processes behind different smile reactions, in different social and emotional contexts, which may overcome possible biases inherent in behavioral and self-report measures.

## 5. Conclusions

Research in embodied cognition has demonstrated that language is bodily grounded and that words are able to get under our skin in ways that are becoming increasingly clear, as they are variegated and complex (Fino et al., 2016, 2019; Foroni & Semin, 2009; Gallese & Lakoff, 2005). However, the extent to which they are able to do so relies on the larger social context and is sensibly modulated by social factors. Detecting highly distinct smile reactions to linguistic portrayals of ingroup and outgroup politicians’ expressions adds another layer to our understanding of how language and social information can shape embodied effects in a highly nuanced fashion.

## Notes

### Competing Interest Statement

The authors have declared no competing interest.

